# Developmental overproduction of cortical superficial neurons impairs adult auditory cortical processing

**DOI:** 10.1101/2022.02.09.479758

**Authors:** Mirna Merkler, Nancy Y Ip, Shuzo Sakata

## Abstract

While evolutionary cortical expansion is thought to underlie the evolution of human cognitive capabilities, excessive developmental expansion can lead to megalencephaly, often found in neurodevelopmental disorders such as autism spectrum disorder. Still, little is known about how the overproduction of cortical neurons during development affects cortical processing and behavior in later life. Here we show that the developmental overproduction of cortical superficial neurons impairs auditory processing in mice. We took advantage of a WNT/β- catenin signaling inhibitor, XAV939, to overproduce cortical superficial excitatory neurons during development. XAV939-treated adult mice exhibited a longer reaction time and a higher threshold to detect acoustic stimuli behaviorally. This mouse model also demonstrated abnormal auditory cortical processing depending on experimental conditions: in a passive listening condition, we observed lower beta power and lower spontaneous and auditory-evoked activity in putative excitatory cells whereas higher pre-stimulus spontaneous activity in excitatory cells is associated with failing to detect auditory stimuli behaviorally. On the other hand, the auditory thalamus did not show any significant difference in neural firing between XAV939-treated and control groups. Furthermore, functional monosynaptic connections were significantly reduced between cortical putative excitatory cells. Altogether, our results suggest that the atypical auditory detectability of XAV939-treated animals can be explained by abnormal auditory cortical processing. Although the expansion of cortical size is evolutionarily advantageous, an abnormal expansion during development can result in detrimental effects on cortical processing and perceptual behavior in adulthood.

## Introduction

The mammalian brain has evolved the neocortex with a prominent six-layered structure (Douglas and Martin, 2004, Harris and Shepherd, 2015, Striedter, 1997). Over a hundred transcriptionally distinct neuron types can be identified across cortical laminae and they exhibit highly diverse functional properties (Yao et al., 2021, Harris et al., 2019, Gouwens et al., 2019, Markram et al., 2004, Brain Initiative Cell Census Network, 2021, Hodge et al., 2019). Although we have witnessed tremendous progress on the characterization of neuronal diversity and circuit operations in the neocortex over the past decade, neuronal number has also long been considered a crucial factor for neural function (Williams and Herrup, 1988, Herculano-Houzel, 2009). For instance, the number of active neurons has been recognized as a fundamental feature to realize efficient neuronal processing (Barlow, 1972, Olshausen and Field, 2004). Nonetheless, we still know little about to what extent the changes in the number of cortical neurons affect neural computation.

The number of cortical neurons is primarily determined by two fundamental factors in development, that is, the length of the neurogenic period and the number of neurons generated per unit of time (Wilsch-Brauninger et al., 2016, Lui et al., 2011). Deficits in neural stem cell proliferation and differentiation can lead to neurodevelopmental disorders, such as autism spectrum disorder (ASD) and intellectual disability (Iakoucheva et al., 2019, Ernst, 2016). While a wide range of anatomical abnormalities has been associated with ASD, macrocephaly can be observed in ∼15% of autistic patients (Hazlett et al., 2017, Piven et al., 1995). However, it remains unclear how such abnormalities in cortical development can lead to abnormalities in cortical information processing and behavior in later life.

Over the past decade, various mouse models of neurodevelopmental disorders have been developed (Kazdoba et al., 2016, Silverman et al., 2010). While some models reflect genetic deficits in human patients, environmental factors during development can also contribute to ASD (Kim et al., 2017, Choi et al., 2016). Yet, it is still challenging to manipulate developmental processes, especially the number of neurons, in a time-limited fashion. Here we take advantage of a WNT/β-catenin signaling inhibitor, XAV939 (Huang et al., 2009). This drug inhibits Tankyrase, which stimulates Axin degradation through the ubiquitin-proteasome pathway (Huang et al., 2009). Axin, a master scaffolding protein for cortical development (Fang et al., 2013, Ye et al., 2015), contains several functional domains to interact with signaling molecules including Tankyrase. Mutations in the *Axin* gene are associated with abnormal brain size (Heisenberg et al., 2001, Kavaslar et al., 2000). Consistently, *in utero* microinjections of XAV939 in mice overproduce only cortical superficial excitatory neurons and lead to autism-like behavioral phenotypes (Fang et al., 2014). This treatment does not alter the production of interneurons, microglia or astrocytes, nor does it affect the number of neurons in the hippocampal formation (Fang et al., 2014). Further, treated mice have more neuronal ensembles in visual cortical layer L2/3 and demonstrate improved visual acuity (Fang and Yuste, 2017). However, it remains to be explored if the observed behavioral changes in the visual system can be seen in other sensory systems and if the overproduction of superficial cortical excitatory neurons affects neural activity across cortical layers.

Since ASD patients often exhibit abnormal sensitivity to acoustic stimuli (Robertson and Baron-Cohen, 2017, O’Connor, 2012), we examine how XAV939 treatment in mice during development leads to changes in auditory function in adulthood. We find XAV939-induced hyposensitivity to acoustic stimuli, with respect to behavioral responses and auditory cortical activity. These deficits are associated with weakened functional connections between putative excitatory neurons, but not auditory thalamic activity, suggesting a cortical origin.

## Results

### Overproduction of cortical superficial neurons by *in utero* XAV939 microinjections

To overproduce cortical superficial neurons, we injected XAV939 into the lateral ventricle of mouse embryos at embryonic day (E) 14.5, when intermediate progenitor cells that will differentiate into L2/3 excitatory neurons are proliferated (**Figure 1A**). To examine the effect of XAV939 microinjections, we performed histological analysis at postnatal day (P) 2 and 21 (**Figures 1B-D**). As reported previously (Fang et al., 2014), we confirmed that cortical superficial layers are significantly expanded in treated animals compared to vehicle controls at P2 (*n*_control_ = 15 animals, *n*_treated_ = 16 animals, *p* < 0.05, *t*-test) (**Figure 1C**) and P21 (*n*_control_ = 13 animals, *n*_treated_ = 14 animals, *p* < 0.001, *t*-test) (**Figure 1D**). Consistent with previous reports (Fang et al., 2013), deeper cortical layers, like L5, have shown a slight, non-significant reduction (*p_P2_* = 0.12, *p_P21_* = 0.14, *t*-test) (**Figures 1B, E and F**).

**Figure 1.**
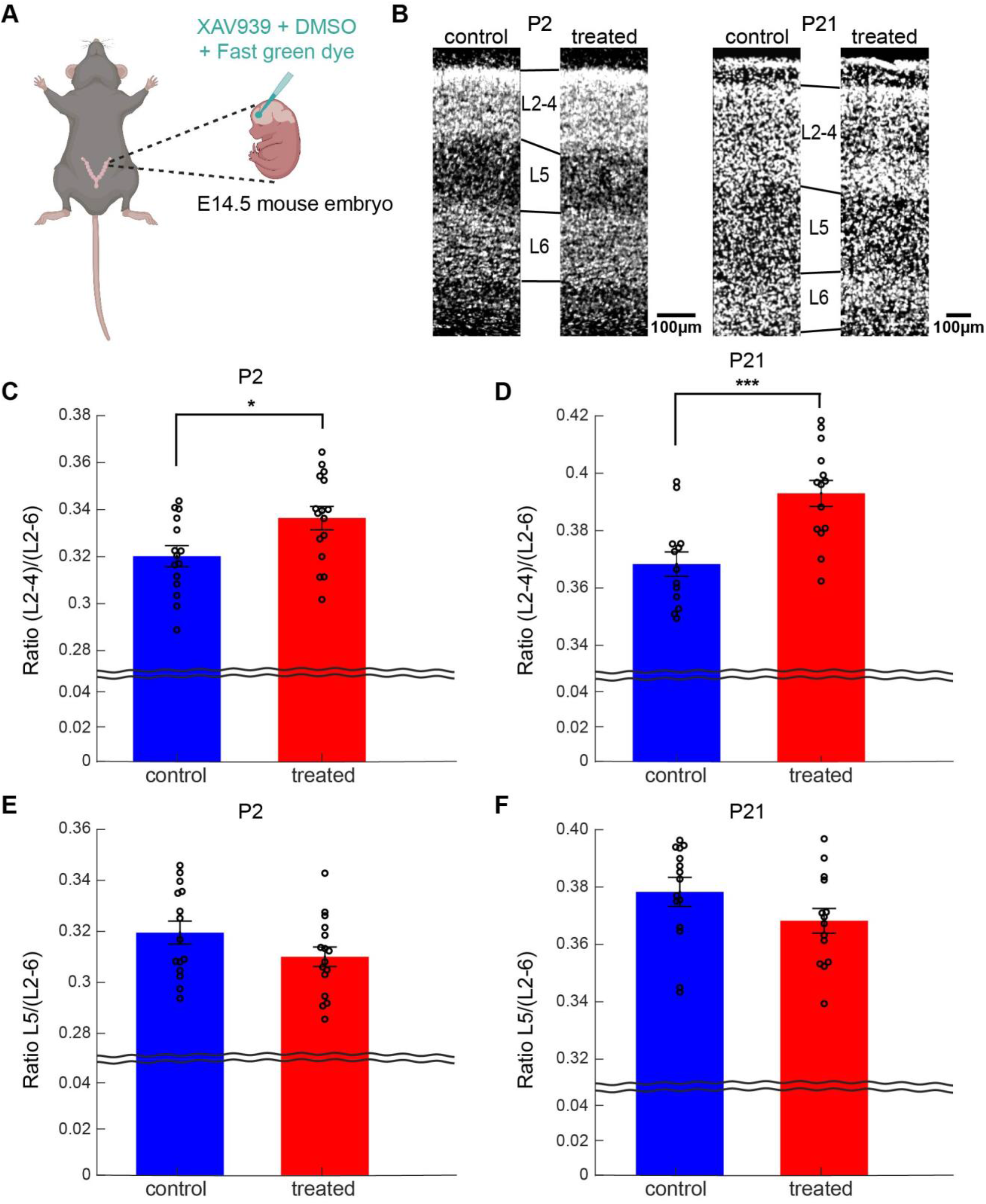
XAV939 caused an increase in the width of cortical superficial layers. **(A)** Schematics of *in utero* microinjections. Image was created with BioRender. **(B)** Comparison of cortical layer width between a control and a treated mouse on postnatal day 2 (P2; left) and P21 (right). **(C and D)** Fraction of cortical superficial layers (L2-4) at P2 **(C)** and P21 **(D)**. **(E and F)** Fraction of L5 at P2 **(E)** and P21 **(F)**. Error bars indicate SEM; \**p* < 0.05, \*\*\**p* < 0.001; *t*-test (**C** and **D**).

Using this animal model, we investigated whether and how the developmental overproduction of cortical superficial neurons affects auditory processing in adults.

### Hyposensitivity to auditory stimulus in XAV939-treated adult mice

To examine the perceptual ability of XAV939-treated mice, we trained adult (∼P70) mice to perform an auditory detection task in a head-restrained condition (**Figure 2**). In this task, mice were required to perform a licking response when they perceived a broadband white noise with varied intensities. When mice correctly detected a sound, they were rewarded with water. In a “catch” trial, no sound was presented. We found a significant effect of treatment on detection performance, measured by *d’* (*n*_control_ = 22 sessions, *n*_treated_ = 32 sessions, *F*_1,53_ = 4.11, *p* < 0.05, two-way ANOVA) (**Figure 2D**). We also estimated detection threshold by applying a logistic regression analysis (see Materials and Methods). We found a significantly higher detection threshold of XAV939-treated mice compared to that of control mice (*n*_control_ = 22 sessions, *n*_treated_ = 32 sessions, *p* < 0.05, rank sum test) (**Figure 2D** inset). Consistent with these observations, the effect of treatment on the average reaction time was also significant (*n*_control_ = 22 sessions, *n*_treated_ = 32 sessions, *F*_1,53_ = 12.87, *p* < 0.0005, two-way ANOVA) (**Figure 2E**). Thus, the developmental overproduction of cortical superficial neurons led to a hyposensitivity to auditory stimulus in adult mice.

**Figure 2.**
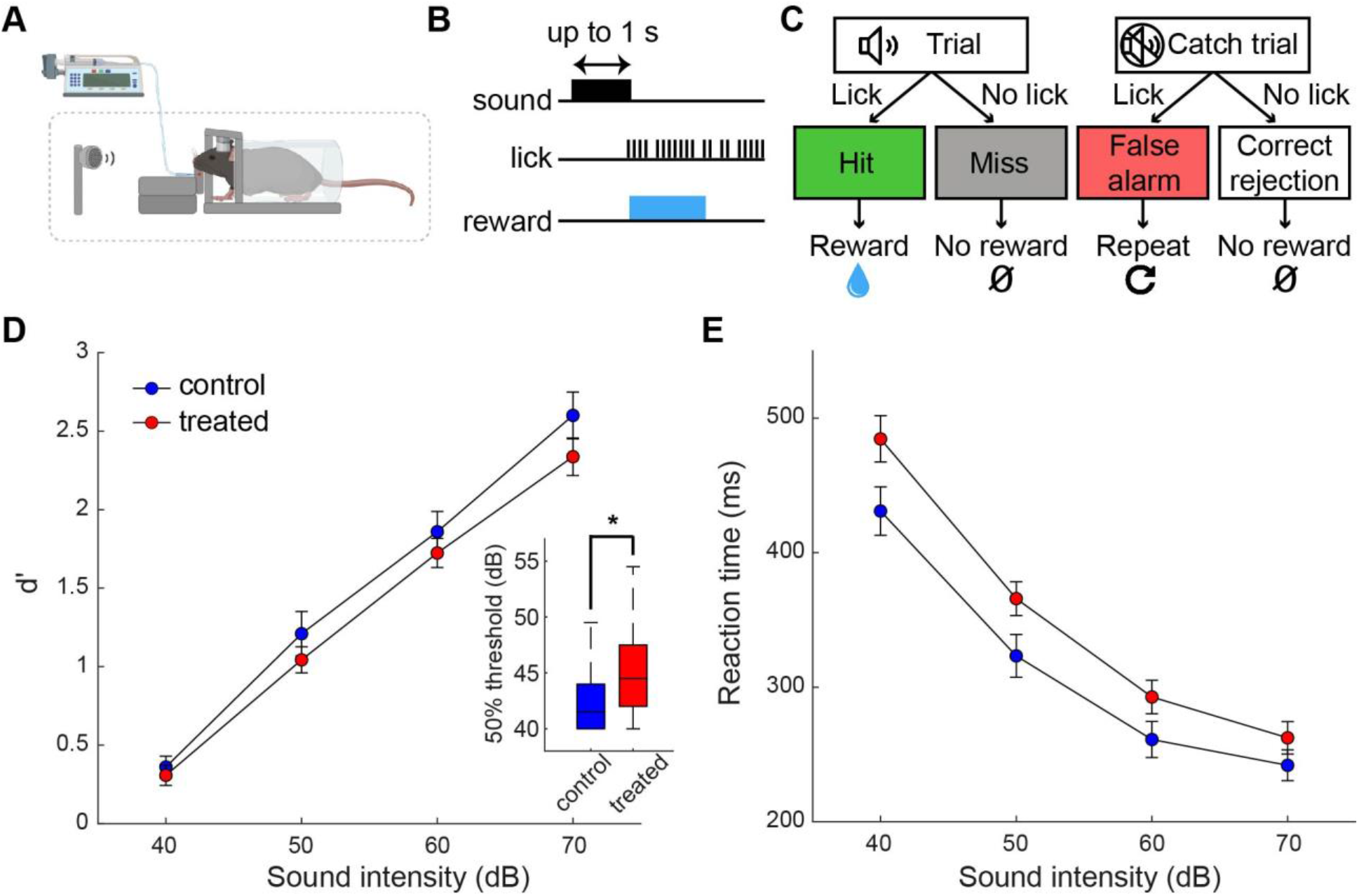
XAV939 treated mice exhibit hearing deficits. **(A)** Schematics of head-fixed behavioral training. Elements located inside a soundproof box are positioned within the dashed box. Image was created with BioRender. **(B)** Timeline of events in the auditory detection task. Trial starts with the sound onset. If the mouse licks during the response window (lasting 1 s), the sound turns off and water reward is delivered. **(C)** Diagram of possible events and outcomes during an auditory detection task session. **(D)** Comparison of *d’* between control and treated mice across sound intensities (*n* = 22 vs. 32 sessions; 2 sessions per mouse). Inset: Detection threshold comparison between control and treated mice. **(E)** Comparison of reaction time between control and treated mice across sound intensities (*n* = 22 vs. 32 sessions; 2 sessions per mouse). Error bars indicate SEM; \**p* < 0.05; Rank sum test (**D** inset), two-way ANOVA (**D** and **E**),

### Abnormal spontaneous population activity in XAV939-treated auditory cortex

To investigate the underlying neural mechanisms of the hyposensitivity to auditory stimulus, we performed *in vivo* electrophysiological recordings in both passive listening and task-performing conditions (**Figures 3-7**). We used either 64-channel silicon probes or Neuropixels probes to monitor population activity (**Figure 3A**): the latter probes allowed monitoring activity from both the auditory cortex and the auditory thalamus simultaneously. In the auditory cortex, we recorded from 463 neurons (382 from 13 passive listening animals across 24 recordings; 81 from 3 task-performing animals across 4 recordings) in treated animals and 171 neurons (123 from 10 passive listening animals across 18 recordings; 48 from 3 task-performing animals across 5 recordings) in control animals. In the auditory thalamus, we recorded from 127 neurons (from 6 passive listening animals across 11 recordings) in treated animals and 62 neurons (from 3 passive listening animals across 6 recordings) in control animals. We further classified cortical neurons into broad-spiking (BS; putative excitatory) neurons (*n*_control_ = 129; *n*_treated_ = 344) and narrow-spiking (NS; putative fast-spiking) neurons (*n*_control_ = 42; *n*_treated_ = 119) based on their spike waveforms (**Figure 3B**).

**Figure 3.**
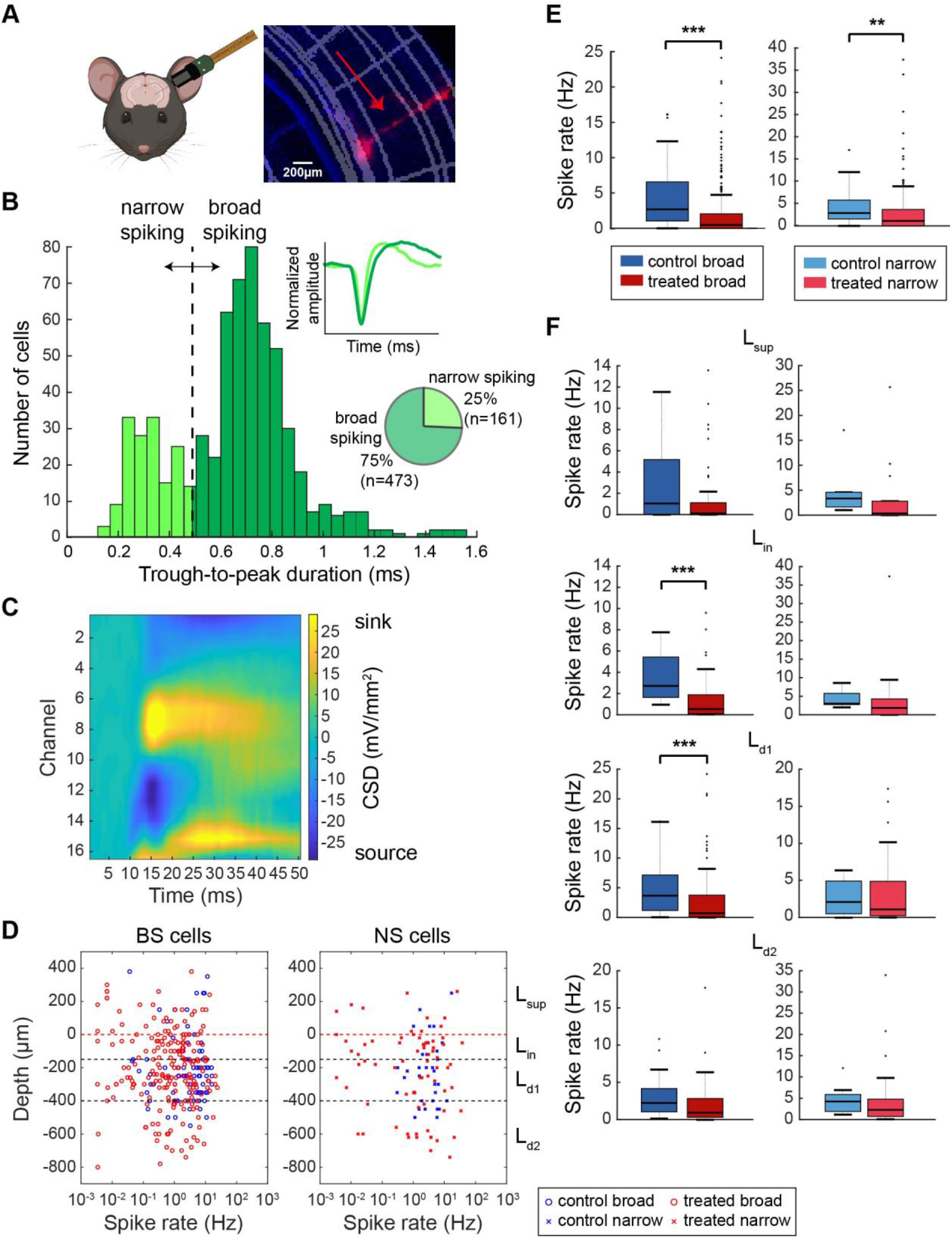
Spontaneous cortical activity was significantly altered in treated mice. **(A)** Schematics of Neuropixels probe insertion (left) and histological confirmation of the probe location in the auditory cortex (right). The schematics was created with BioRender. **(B)** Cell type classification depending on their waveform trough to peak duration. Two clusters (broad-spiking; BS and narrow-spiking; NS) were split at 0.5 ms border. Top inset: comparison between narrow and broad waveform. Bottom inset: Total number and percentages of BS and NS neurons across all recordings (passive listening and task performing), used in further analysis. **(C)** Current source density (CSD) example for one shank (16 channels) of a silicon probe recording. Top current sink channel (yellow; channel 8) is set as depth zero and the top border of input layer (L_in_). Using CSD, neurons were divided into cortical layers depending on the depth of the channel with their largest amplitude of average spike waveforms. **(D)** Scatter plots of BS (left) and NS (right) neurons, showing their depth and cortical layer position as well as their spontaneous activity. Estimated layer borders are as follows: Superficial layers (L_sup_) > 0 µm, input layer (L_in_) 0 to -150 µm, deep layer 1 (L_d1_) -150 to -400 µm, deep layer 2 (L_d2_) < -400 µm. **(E)** Spontaneous activity in control and treated mice (*n* = 89 vs. 281 BS cells, and 34 vs. 101 NS cells). **(F)** Spontaneous activity across cortical layers. L_sup_, putative L2/3; L_in_, putative L4; L_d1_, putative L5; L_d2_, putative L6. \*\**p* < 0.01, \*\*\**p* < 0.001; Rank sum test (**E** and **F**).

Analyzing these datasets, we firstly examined whether and how spontaneous activity in the auditory cortex was modified in XAV939-treated animals under a passive listening condition (**Figures 3E and F**). We found that the spontaneous firing rate of BS (*n*_control_ = 89 cells, *n*_treated_ = 281 cells, *p* < 0.001, rank sum test) (**Figure 3E, left**), as well as NS neurons (*n*_control_ = 34 cells, *n*_treated_ = 101 cells, *p* < 0.01, rank sum test) (**Figure 3E, right**) was significantly lower in treated animals compared to that in control animals. Based on the depth profile of spike waveforms and current source density (CSD), we broke down auditory cortical neurons according to their depths into superficial layers (L_sup_; putative L2/3), input layer (L_in_; putative L4), deep layer 1 (L_d1_; putative L5) and deep layer 2 (L_d2_; putative L6) (**Figures 3C and D**, see also Materials and Methods). Across layers, we observed lower firing rate of BS neurons in treated mice, with significant difference in L_in_ and L_d1_ (*p* < 0.001, rank sum test) (**Figure 3F**).

We also assessed spontaneous neural oscillations between groups based on local field potentials (LFPs) (**Figures 4A and B**). The power in higher frequency bands (specifically, alpha and beta) was reduced in treated animals (*p*_alpha_ = 0.08, *p*_beta_ < 0.05; *n*_control_ = 17 recordings, *n*_treated_ = 19 recordings, *t*-test), while delta power was slightly, but non-significantly, enhanced (*p*_delta_ = 0.11). In **Figure 4B**, we also assessed LFP powers across cortical layers and frequency bands. The effect size (Hedge’s *g*) was consistently high (∼1) across layers in beta band, along with low (∼-1) and high (>1) effect size for deep layers in the delta and alpha bands, respectively.

**Figure 4.**
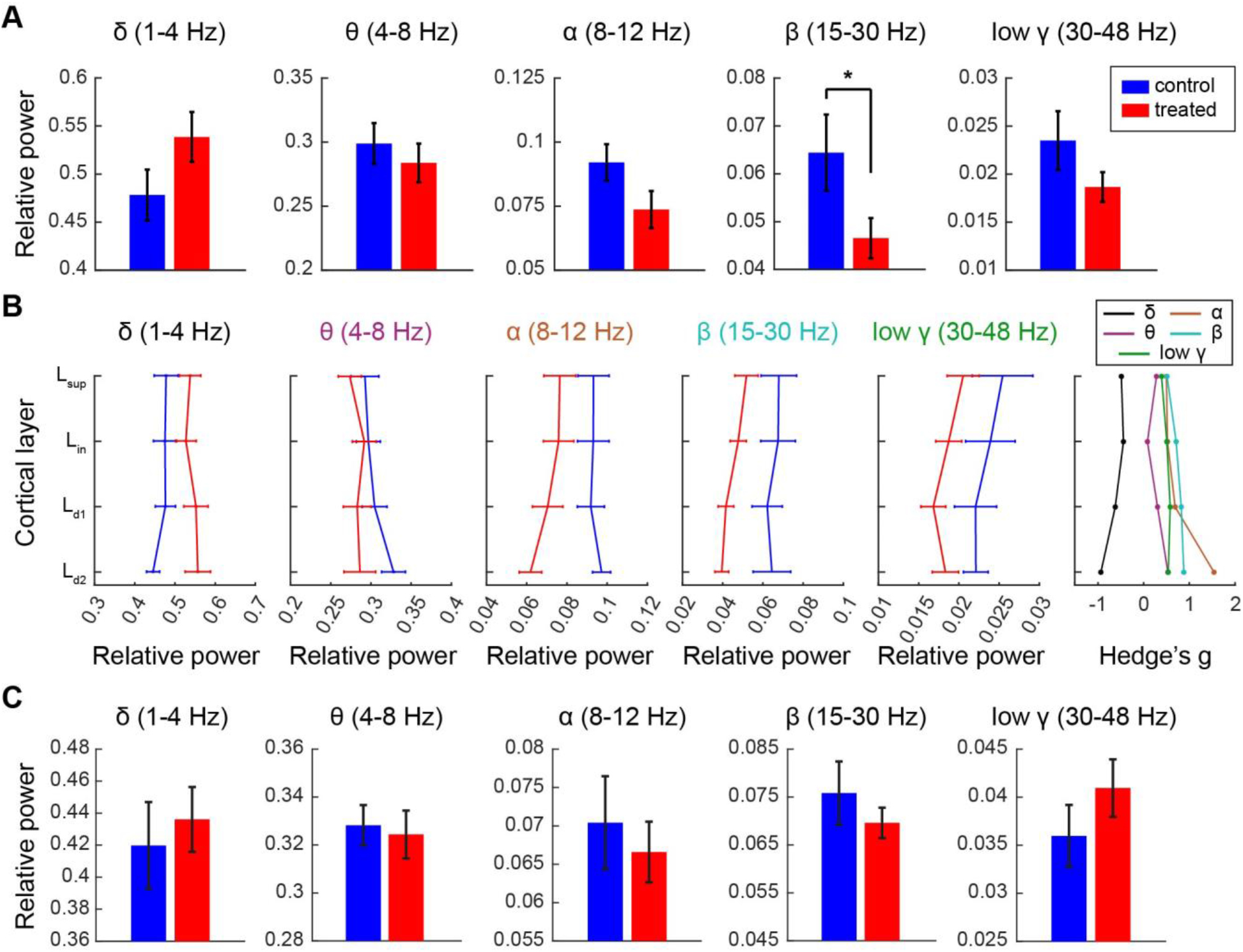
Local field potential changes in treated mice were confined to the cortex. **(A)** Comparison of relative power across frequency bands in the auditory cortex of control and treated mice during silence. Mean relative power was calculated across cortical channels for each recording (*n* = 17 vs. 18 recordings). **(B)** Relative resting state power across cortical layers, for each frequency band. Hedge’s *g* was calculated to look at the effect of treatment for each cortical layer across frequency bands. **(C)** Mean relative power across hippocampal channels during silence, for each frequency band (n = 6 vs. 16 recordings). Error bars indicate SEM; \**p* < 0.05; *t*-test (**A** and **C**).

By taking the advantage of Neuropixels probe recording, we also assessed simultaneously monitored LFPs in the hippocampus to examine if this difference might originate from other regions (**Figure 4C**). We did not find any significant differences in oscillatory powers across frequency bands between two animal groups (*n*_control_ = 6 recordings, *n*_treated_ = 16 recordings, *p*_delta_ = 0.67, *p*_theta_ = 0.83, *p*_alpha_ = 0.61, *p*_beta_ = 0.35, *p*_gamma_ = 0.36; *t*-test). These results indicate that the overall spiking activity of BS neurons reduces, and cortical oscillatory power becomes lower in XAV939-treated animals.

### Abnormal auditory-evoked activity in XAV939-treated auditory cortex

Because the behavioral detection threshold was significantly elevated in the treated mice (**Figure 2D**), we hypothesized that auditory evoked responses in the auditory cortex of the treated animals are diminished. To test this, we began by comparing auditory evoked responses in BS and NS cells in treated and control mice (**Figures 5A and B**). As the sound intensity increased, evoked responses became larger. However, we noticed that evoked responses of BS neurons in treated mice were consistently lower than those in control mice (**Figure 5A**). To confirm this quantitatively, we computed the mean spike rate in a 50 ms window from the stimulus onset across cells (**Figures 5C and D**). In BS cells, auditory-evoked responses were significantly reduced in treated animals across all sound intensities (*n*_control_ = 89 cells, *n*_treated_ = 281 cells, *p* < 0.001, rank sum test with Bonferroni correction) (**Figure 5C**). NS neurons in treated mice tended to exhibit slightly lower auditory-evoked responses, but with no significance (*n*_control_ = 34 cells, *n*_treated_ = 101 cells, *p* > 0.05 across all intensities, rank sum test with Bonferroni correction) (**Figure 5D)**.

**Figure 5.**
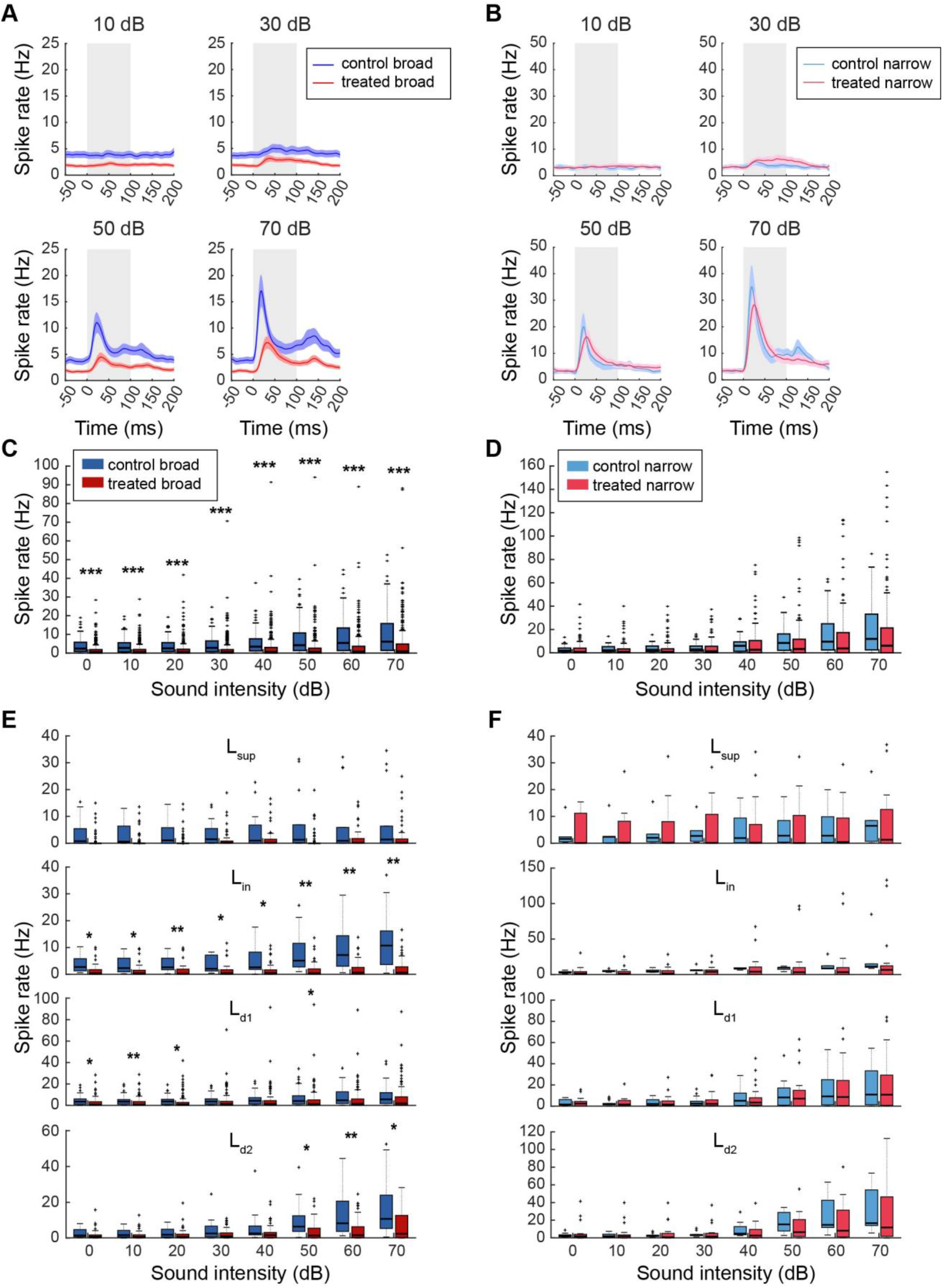
Auditory evoked activity was affected by XAV939 treatment. **(A and B)** Peristimulus time histogram (PSTH) of activity (mean ± SEM) of BS neurons **(A)** and NS neurons **(B)** in control and treated group during sound presentation. Sound period is labelled as a gray background (lasting 100ms). **(C and D)** Spike rate of BS **(C)** and NS **(D)** neurons during first 50 ms of sound presentation. **(E and F)** Activity of cells shown in **(C)** and **(D)** split into cortical layers. \**p* < 0.05, \*\**p* < 0.01, \*\*\**p* < 0.001; Rank sum test with Bonferroni correction (**C – F**).

We further assessed the laminar specificity in firing rate changes (**Figures 5E and F**). Evoked responses tended to be lower across layers, with significant reduction in L_in_, L_d1 and_ L_d2_ BS neurons. On the other hand, NS cells exhibited diverse effects (**Figure 5F**). Overall, XAV939-treatment led to a reduction in auditory evoked responses in BS neurons, but not NS neurons, across cortical layers.

### Abnormal auditory cortical activity in task-performing XAV939-treated mice

Next, we directly compared auditory cortical activity in task-performing animals between the two groups (**Figure 6A**). As neural activity in a pre-stimulus period can influence behavioral performance (Schölvinck et al., 2012), we examined if the spontaneous activity in the pre-stimulus window differed between animal groups depending on their behavioral outcomes, i.e., “hit” and “miss”. In hit trials, we found significantly lower spontaneous activity of BS neurons in XAV939-treated mice compared to control mice (*n*_control_ = 40, *n*_treated_ = 63, *p* < 0.05, rank sum test) (**Figure 6B, left**). On the other hand, spontaneous activity in miss trials was not significantly different between the groups (*n*_control_ = 40, *n*_treated_ = 63, *p* = 0.15, rank sum test) (**Figure 6B, left**). Although the spontaneous activity is similar between hit and miss trials in control mice (*p* = 0.56, rank sum test), the spontaneous activity in treated mice was significantly higher in miss trials compared to hit trials (*p* < 0.01, rank sum test). Despite these differences in BS cells across conditions, we did not observe significant changes in spontaneous activity of NS neurons regardless of the outcome (*n*_control_ = 8, *n*_treated_ = 18, *p*_hit_ = 0.58, *p*_miss_ = 0.14, rank sum test) (**Figure 6B, right**).

**Figure 6.**
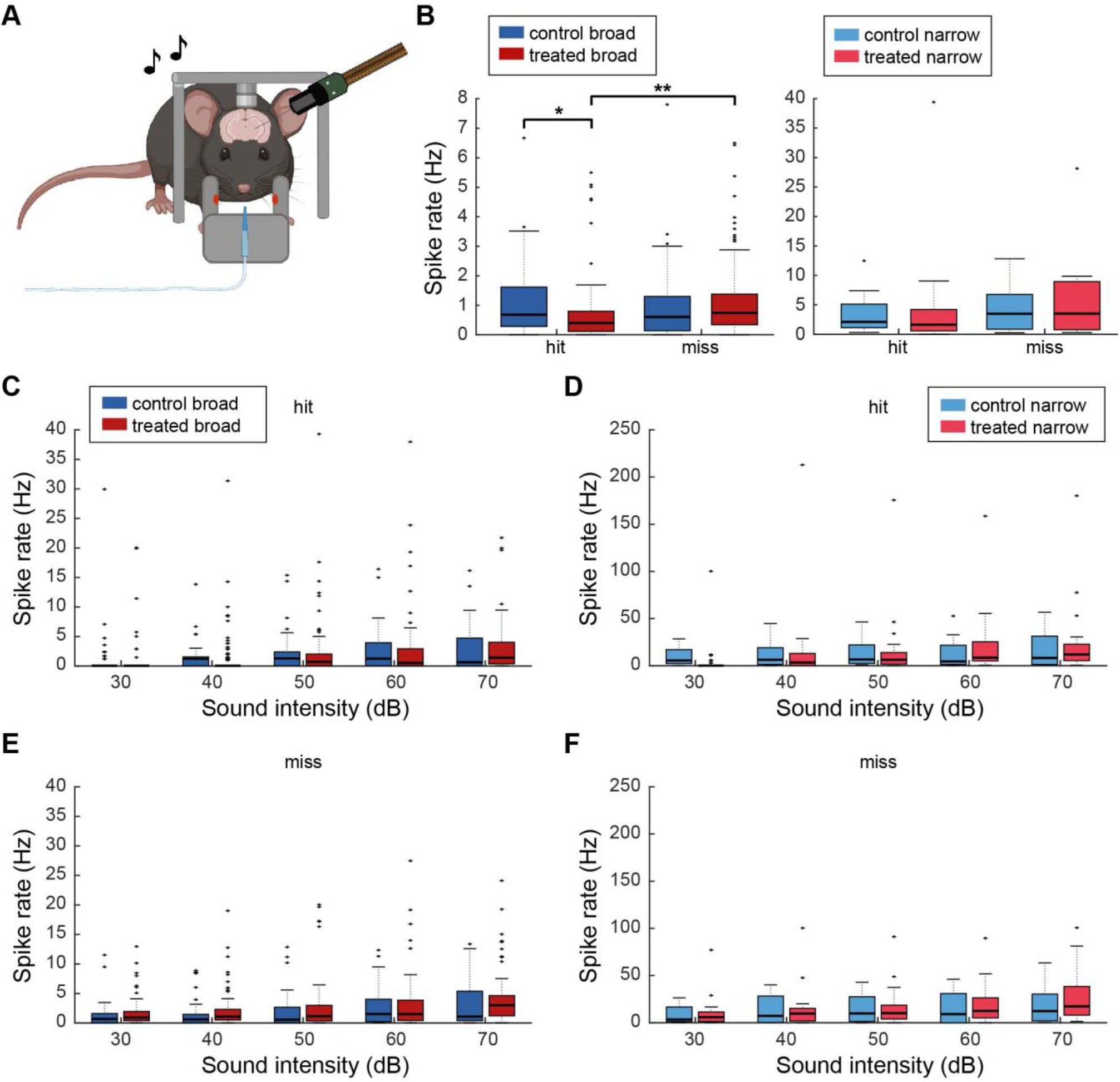
Trial outcome related differences in neuronal activity were observed in task behaving treated mice. **(A)** Schematics of Neuropixels recording in task behaving head-fixed conditions. Image was created with BioRender. **(B)** Spontaneous activity during prestimulus window (0.5 s before trial onset), before trials with “hit” or “miss” outcome (n = 40 vs. 63 BS cells, and 8 vs. 18 NS cells). **(C and D)** Activity of BS **(C)** and NS **(D)** neurons in trials with “hit” outcome, during first 50 ms from sound onset. **(E and F)** Activity of BS **(E)** and NS **(F)** neurons in trials with “miss” outcome. Error bars indicate SEM; \**p* < 0.05, \*\**p* < 0.01; Rank sum test (**B**), rank sum test with Bonferroni correction (**C-F**).

We also looked at the auditory evoked responses in each group for hit and miss trials. The activity of BS neurons of treated mice in hit trials tended to be lower (**Figure 6C**), especially for lower sound intensities, whereas this trend was absent in miss trials (**Figures 6E**). However, the differences between groups in hit trials were not significant (*n*_control_ = 40, *n*_treated_ = 63, *p* > 0.05, across all intensities, rank sum test with Bonferroni correction). NS neurons did not exhibit any significant changes or trends in activity (*n*_control_ = 8, *n*_treated_ = 18, *p* > 0.05, across all intensities, rank sum test with Bonferroni correction) (**Figures 6D and F**). Thus, the poor detection performance of XAV939-treated mice was associated with a bigger change in pre-stimulus activity of BS neurons, rather than auditory evoked activity.

### Auditory thalamic activity in XAV939-treated mice

To examine whether the observed abnormal activity in the auditory cortex of XAV939-treated animals is inherited from an upstream structure, we analyzed activity in the auditory thalamus (**Figure 7**). Neurons were recorded from multiple auditory thalamic nuclei, not just the ventral division of the medial geniculate body (MGBv) (**Figure 7B**). We found that MGB neurons did not show any significant differences in spontaneous (*n*_control_ = 62, *n*_treated_ = 127, *p* = 0.14, rank sum test) (**Figure 7C**) or auditory evoked activity between two animal groups (*n*_control_ = 62, *n*_treated_ = 127, *p* > 0.3 across all intensities, rank sum test with Bonferroni correction) (**Figure 7D**). These results imply cortical mechanisms for the hypofunction of auditory processing in XAV939-treated mice.

**Figure 7.**
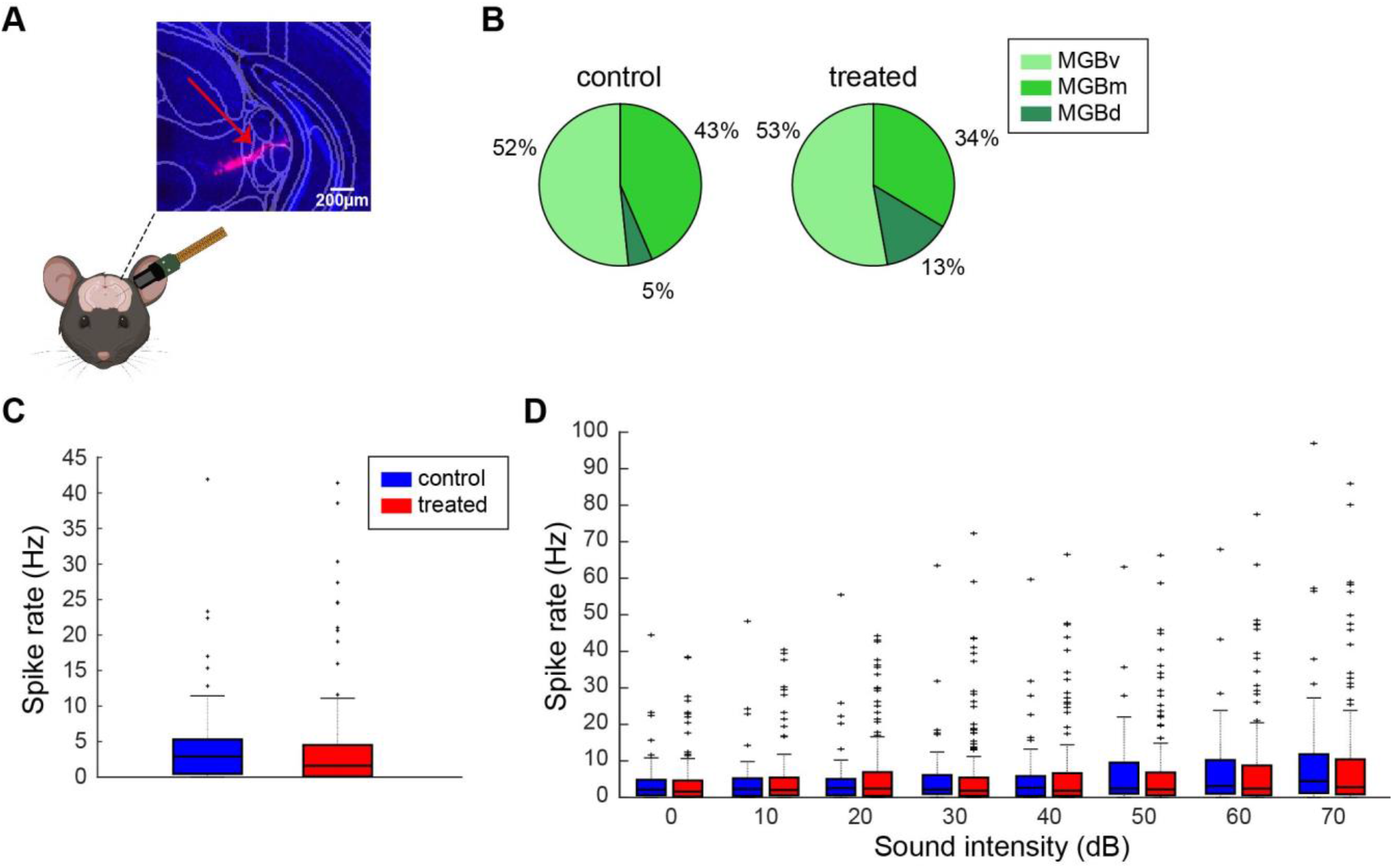
Auditory thalamic activity was not affected by XAV939 treatment. **(A)** Schematics of simultaneous Neuropixels recording from auditory cortex (AC) and medial geniculate body (MGB), with histological confirmation of probe location in MGB. **(B)** Percentage of MGB neurons recorded across different nuclei in control (left) and treated (right) mice. In our recordings, most of the neurons came from MGBv, with only a small fraction of MGBd neurons. **(C)** Spontaneous activity in MGB during passive listening conditions (*n* = 62 vs. 127 cells). **(D)** MGB activity in passive listening conditions across sound intensities. None were statistically significant (*p* > 0.1); Rank sum test **(C)**, rank sum test with Bonferroni correction **(D)**.

### Reduced effective excitatory connections in XAV939-treated auditory cortex

To explore a potential mechanism to explain abnormal auditory cortical processing in XAV939-treated animals, we inferred monosynaptic connections among simultaneously recorded neurons by computing cross-correlograms (CCGs) (**Figures 8**). This approach allows us to identify strong synaptic coupling between neurons based on their spike trains (Fujisawa et al., 2008). As shown in **Figure 8A**, there are three types. Firstly, if there is no strong coupling, no significant modulation can be seen. On the other hand, if there are strong effective connections from one neuron to another, significant excitatory or inhibitory modulations can be seen within several milliseconds after spikes. We systematically assessed these effective couplings across all simultaneously recorded neurons in the auditory cortex (**Figures 8B-D**). Across datasets, such connections were generally sparse in both groups (**Figure 8B**). However, when we compared the connection probability of individual neurons between two animal groups, we found significantly lower connection probabilities in XAV939-treated mice from BS to BS neurons (*n*_control_ = 129, *n*_treated_ = 340, *p* < 0.01, rank sum test with Bonferroni correction) (**Figure 8C**). We also noticed a trend toward a reduced connection probability coming from NS neurons, although this was not statistically significant (*n*_control_ = 45, *n*_treated_ = 120, *p* > 0.05, rank sum test with Bonferroni correction) (**Figure 8D**). These results suggest weakened cortical synaptic connections in XAV939-treated mice.

**Figure 8.**
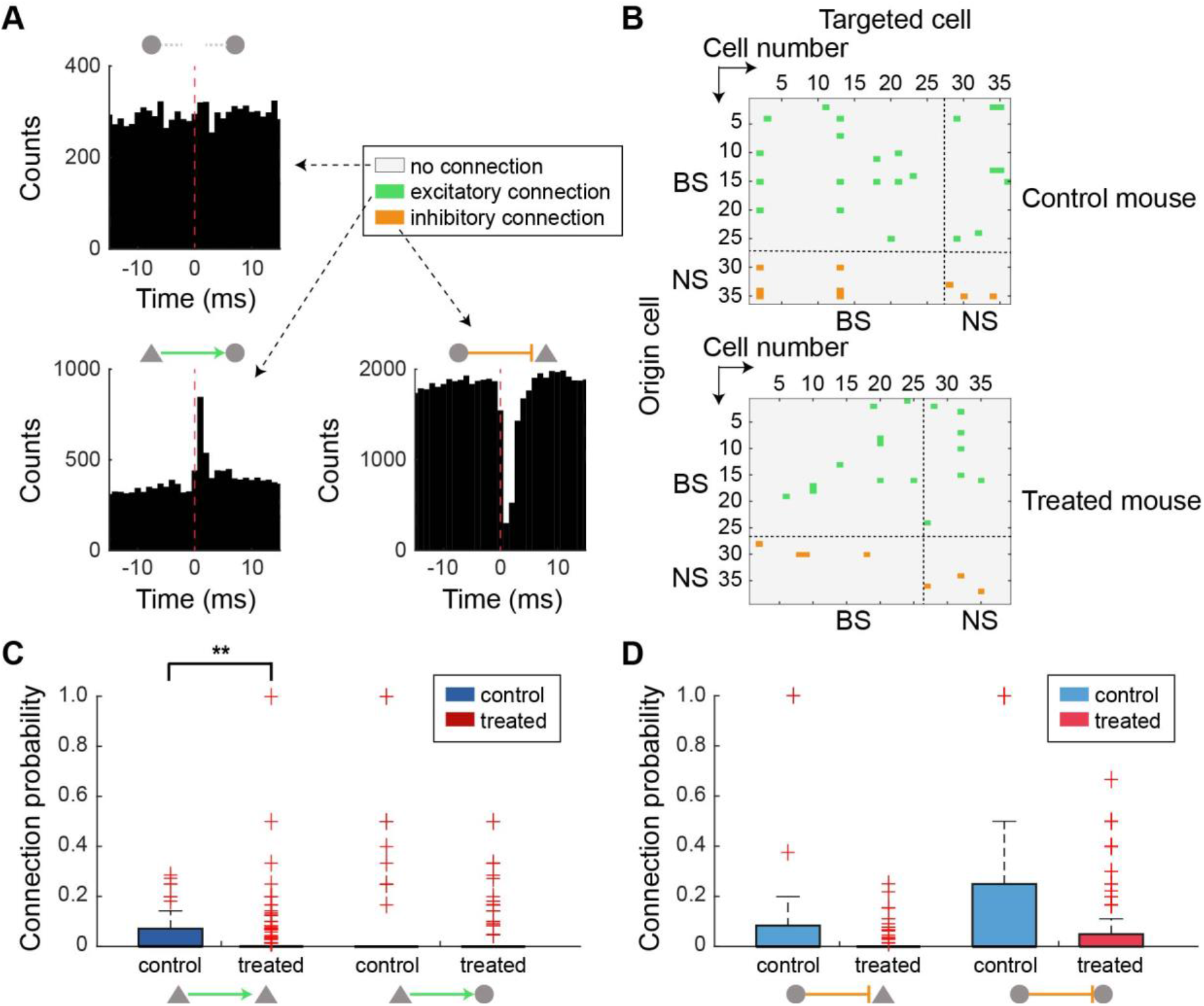
XAV939 treated mice have fewer monosynaptic connections within the auditory cortex. **(A)** Examples of cross-correlograms between neuron pairs; neurons that are not connected (top), excitatory connection (bottom left) and inhibitory connection (bottom right). Red dotted line marks the spike time of origin cell, while histogram shows the activity of targeted cell. **(B)** Connectivity matrices from one control and one treated mouse, showing the connections between simultaneously recorded neuron pairs. **(C)** Probability of an excitatory connection coming from BS neurons (triangle; *n* = 129 vs. 340) to either BS (triangle) or NS neurons (circle). **(D)** Probability of an inhibitory connection coming from NS neurons (circle; *n* = 42 vs. 120). \*\**p* < 0.01; Rank sum test with Bonferroni correction (**C** and **D**).

## Discussion

We examined how manipulating the number of cortical neurons during development affects cortical processing in adulthood. We found that XAV939-treated mice exhibit a reduction in behavioral performance during the auditory detection task, as well as having reduced spontaneous activity in BS neurons of the auditory cortex. In addition, the putative monosynaptic excitatory connections between BS neurons were weakened. On the other hand, the auditory thalamus did not exhibit abnormal activity in XAV939-treated mice. Altogether, these results indicate that developmental overproduction of cortical superficial excitatory neurons leads to the hypofunction of auditory processing, possibly due to weakened cortical excitatory synaptic connections.

Although XAV939 has been used to increase cortical superficial neurons previously, effects of XAV939 microinjections on adult behaviors are varied (Fang et al., 2014, Fang and Yuste, 2017): XAV939-treated mice exhibited autistic behaviors, such as repetitive behaviors and social deficits (Fang et al., 2014), and their visual discrimination ability was enhanced (Fang and Yuste, 2017). Unexpectedly, we observed a reduced detection ability in the auditory system. This contradiction can be explained by several factors. Firstly, it is possible that different modalities may exhibit distinct effects. Secondly, the overproduction of cortical superficial neurons may lead to distinct impacts on perceptual detection and discrimination. Because superficial excitatory populations encode sensory information sparsely (de Kock et al., 2007, Crochet et al., 2011, Sakata and Harris, 2009), the overproduction of superficial excitatory neurons may increase a coding space, leading to a better discrimination ability whereas this may not be the case for perceptual detection. Indeed, some ASD subjects exhibit decreased signal detection ability while others report enhanced discrimination of simple stimuli in the visual domain (Bertone et al., 2005, Sanchez-Marin and Padilla-Medina, 2008). In addition, this apparent contradiction may be analogous to the fact that ASD patients often exhibit superior performance for details of their perceptual world, but this comes at a cost for the global perceptive (Robertson and Baron-Cohen, 2017, Shah and Frith, 1983). Future studies need to reconcile this discrepancy.

Why did the overproduction of superficial excitatory neurons diminish the overall cortical activity? We hypothesize that mechanisms have emerged to compensate for the enhanced excitability in cortical circuits during postnatal development. It is consistent with the previous finding that XAV939 administration shifts the balance between excitatory and inhibitory synapses (Fang et al., 2014): the ratio of excitatory to inhibitory synapses in L2/3 in XAV939-treated animals increased at P18, whereas it decreased at P60. This change was associated with a reduction in field excitatory postsynaptic potentials, which is consistent with the present results. Thus, the excitation-inhibition balance may be changed over a period of time in the XAV939 model. During postnatal development, programmed cell death plays a prominent role in establishing appropriate numbers of excitatory and inhibitory cortical neurons (Verney et al., 2000, Southwell et al., 2012). Moreover, accumulating evidence has also demonstrated postnatal activity-dependent processes which sculpt cortical circuits (Thion et al., 2019, Wong et al., 2018). Thus, the overproduction of cortical superficial excitatory neurons during the prenatal period can trigger various compensatory mechanisms, resulting in reduced cortical excitability in adults.

We have noticed a shift in the power of spontaneous LFPs for specific frequency bands in XAV939-treated mice. Our mouse model showed a significant decrease in beta, and a slight decrease in alpha. Alpha band was proposed to stem from rhythmic GABAergic activity (Klimesch, 2012), and beta power increases with GABAergic inhibition (Jensen et al., 2005). Since cortical oscillations represent transitions between phases of low and high excitability (Bishop, 1932, Schroeder and Lakatos, 2009), reduced cortical inhibitory and excitatory spontaneous activity, as well as the functional connectivity differences observed in XAV939 mice could be the source of long-term changes in the oscillatory power across these frequency bands.

As in passive listening conditions, we also observed significantly reduced spontaneous activity of BS neurons before “hit” trials in behaving XAV939-treated animals. Additionally, in treated mice, spontaneous activity before miss trials was significantly higher than before hit. On the other hand, we did not observe any significant differences in evoked responses between animal groups. These results suggest that contrary to the passive listening condition, higher background activity in XAV939-treated mice may lead to behavioral hyposensitivity to acoustic stimuli.

Although the expansion of cortical size is advantageous on the evolutionary timescale (Herculano-Houzel, 2009), our results are consistent with the notion that abnormal expansion of cortical layers during development can lead to an imbalance of excitation and inhibition in the cortical network, resulting in detrimental effects, which can be seen in various neurodevelopmental disorders (Tang et al., 2021, Nelson and Valakh, 2015, Marin, 2012). Indeed, increased cortical volume and L2 neuron number have been reported in ASD (Falcone et al., 2021, Courchesne et al., 2011, Hazlett et al., 2017), and some ASD patients experience hypofunction of sensory systems (Robertson and Baron-Cohen, 2017). Such hypofunction may stem from altered cortical connections, and reestablishing synaptic connections may be a promising therapeutic strategy for ASD.

In conclusion, the overproduction of cortical superficial excitatory neurons during development weakens synaptic connections within BS cortical population in adulthood. We believe that these altered synaptic connections could lead to the observed decrease in spontaneous and evoked BS population activity in our treated mice, as well as a higher behavioral threshold to detect auditory signals.

## Acknowledgements

This work was supported by the BBSRC (BB/M009050X/1), MRC (MR/V033964/1) and Horizon2020-ICT (DEEPER, 101016787) to SS, and the Area of Excellence Scheme of the University Grants Committee (AoE/M-604/16) to NYI.

## Author contributions

MM and SS designed and conceived the project and analyzed the data. MM performed all experiments. SS supervised MM. NYI provided technical support for in utero microinjections. MM and SS wrote the manuscript with input from NYI.

## Materials and Methods

### Animal maintenance and XAV939 administration

All animal experiments and procedures were performed in accordance with the United Kingdom Animals (Scientific Procedures) Act of 1986 Home Office regulations and approved by the Home Office (PPL70/8883 and PP0688944) and the University of Strathclyde’s Ethical Committee.

Pregnant female wild-type C57BL/6 mice (*n* = 32) were housed in a 12 h light/dark cycle. *In utero* microinjections were performed on embryonic day 14.5 (E14.5), where embryos were injected with 0.15 or 0.4 μl of 10 μM XAV939 (X3004, Sigma Aldrich) in DMSO solution with 0.05% Fast Green dye (F7258, Sigma Aldrich; treated litter) or DMSO with 0.05% Fast Green dye solution (control litter), in the lateral ventricle. Mice were perfused on postnatal day 2 (P2) (*n* = 31) and P21 (*n* = 27) for histological assessment of cortical layers width. Only male mice were kept until adulthood (perfused ∼P120) and used for behavioral (*n* = 30) and electrophysiological (*n* = 33) assessments.

### Surgical procedure for experiments in head-fixed condition

The procedure was done as described elsewhere (Lyngholm and Sakata, 2019, Yague et al., 2017). Briefly, ∼8 week old male mice were anesthetized with isoflurane (1 – 1.5%). To provide analgesia, Lidocaine (2%, 0.1 ml) was administered subcutaneously at the site of the incision and Carprofen (Rimadyl, 10 ml/kg, 0.01% diluted in water for injection) was administered subcutaneously in the back. Five screws (418-7123, RS Components) were implanted in the skull, two in the front (AP +1.5 mm, ML ±1.5 mm), two on the cerebellum (AP -6 mm, ML ±2 mm) and one over the right hemisphere (AP -3 mm, ML 3 mm). The posterior left screw was used as a ground, while the anterior right screw was used for cortical electroencephalogram (EEG) recording. The other screws were implanted for anchoring and stabilization of the head cap. A head-post made of 2 nuts was affixed onto the two frontal skull screws. During this surgery, the location of the future craniotomy site above the left hemisphere was also labelled (2 × 2 mm^2^ at AP -2 mm and ML 4 mm). Dental cement was used to cover the skull screws and surface as well as to secure the head-post in place. After the surgery, mice were single housed in high-roofed cages with ad libitum access to food and water and left to recover for at least 5 days.

### Behavioral experiments

#### Apparatus

A custom-made behavioral apparatus was located in a soundproof box (Med Associates Inc). A head-fixed mouse was placed in a restraining tube, with a spout, made from a pipette tip, in front of his mouth. The tip of the spout was aligned with a sensor (PMU24, Panasonic). The spout was connected by tubing to a 10 ml syringe filled with water and secured in a single-syringe infusion pump (AL-1000, World Precision Instruments). The pump was located outside of the box. To present the sounds, a calibrated speaker (ES1, Tucker-Davis Technologies) was placed ∼15 cm in front of the mouse. Water delivery, sound presentation and sensor information were relayed through NI-DAQ (USB-6343, National Instruments) and controlled by a custom-written LabVIEW program (National Instruments). Broadband white noise was generated digitally (USB-6343, National Instruments) and amplified (System 3 ED1, Tucker-Davis Technologies) before being transmitted to the speaker.

#### Behavioral training

After recovery from surgical procedure, mice were placed into the reverse light/dark cycle and water restriction was initiated a week after the adaptation to the new cycle. The restriction was daily, starting gradually until reaching 0.04 ml/g/day. During the restriction acclimation period of 7 days, mice were also habituated to handling and head fixation. Daily training sessions lasted 1 hr, during which mice received water through a water pump as a reward (1 – 3 µl water drop per pump). Mice were weighed after each session and the remainder of their daily water amount was delivered in the form of hydrogel (70-01-5022, ClearH2O). Behavioral training consisted of 3 phases: basic lick training, auditory conditioning and auditory detection.

In the first training phase (Basic lick training), mice were familiarized with the system. Whenever the mouse licked the spout, their tongue would cross the beam of the sensor activating the pump and water would be delivered. After making more than 300 licks in a session, the animal would move on to the next phase.

In the second phase (Auditory conditioning), mice were trained to lick only when a broadband white noise (60 dB SPL) was present. At first, the sound was presented for 8 s, followed by 2 s of silence. Mice could get a water reward only if they licked during the sound period. After 50 valid trials, the sound would be reduced by 1 s and silence increased by 1 s. The sound presentation was reduced in this way across sessions until reaching 2 s. If an animal had ≥ 150 licks with ≥ 60% success rate for 2 s sound periods in 2 consecutive sessions, they could move on to the final phase.

In the final phase (Auditory detection task), mice were trained and then later assessed in licking whenever they heard a sound, across different intensities. A broadband white noise (30 – 70 dB SPL, 10 dB SPL steps) was presented for a maximum of 1 s. Intertrial interval (ITI) in this phase lasted 2 – 4 s, and both intensities and ITI length were pseudo-randomized across trials. Catch trials were also present in this phase, allowing us to assess how often the mouse engaged randomly. When the difference between the catch trial false alarm rate and 70 dB hit rate was more than 40% in two consecutive sessions, after at least 10 training sessions, the mouse would finish training. These two sessions were used in analysis of behavioral performance.

### In vivo electrophysiology

Detailed recording procedures have been previously described (Lyngholm and Sakata, 2019, Yague et al., 2017). One day before the recording, mice were briefly anesthetized with isoflurane and a craniotomy was done over the auditory cortex (AC) (see above for the coordinate), on the site marked previously during the head cap surgery. Craniotomy site was covered with a biocompatible sealant (Kwik-Sil, World Precision Instruments). During the recording sessions, Kwik-Sil was removed and phosphate buffer saline (PBS) was used to prevent the brain surface from drying. Neural population activity was recorded in ∼1 hr daily sessions over 2 days. Head-fixed, awake mice were sat in a restraining tube within a soundproof box (Industrial Acoustics Company). Electrophysiological recordings were done with either a 64-channel silicon probe (A4×16-6mm-50-200-177-A64, NeuroNexus Technologies; recorded from AC) or with a Neuropixels 1.0 probe (recorded from AC and the medial geniculate body (MGB)), mounted on a manipulator (SM-15 or DMA-1511, Narishige). Probes were inserted perpendicular to the cortical surface (AC recording depth 750-900 µm; MGB recording depth ∼4000 µm). Signals collected through the silicon probe were amplified and digitized (RHD2132 and RHD2000, Intan Technologies Inc.) and recorded using a custom-written LabVIEW program (National Instruments). Signals collected through the Neuropixels probe were amplified and digitized in the probes integrated circuit and recorded using SpikeGLX (Janelia Research Campus). The recording sessions were typically initiated ∼0.5-1 hr after the probe insertion, to allow for signal stabilization. Probes were labelled with either DiI (D-282, Invitrogen, 10% diluted in ethanol) or CM-DiI (C7001, Invitrogen, 0.1% w/v diluted in ethanol) before insertion, thus allowing us to find the probe track later in histology.

#### Sound presentation and behavioral task during electrophysiological recording

For passive listening experiments, broadband white noise (100 – 200 repetitions of 100 ms pulses with 1 ms cosine ramps, 10 dB SPL steps, 0 – 70 dB SPL intensity range presented pseudo-randomly, 500 ms interstimulus interval (ISI)), was generated digitally (sampling rate 97.7 kHz, RZ6 Multi I/O processor, Tucker-Davis Technologies) and transmitted in free-field through a calibrated speaker (ES1, Tucker-Davis Technologies). In behavioral task recordings, broadband white noise was presented for a maximum of 1 s (or until the animal licking), in 30 – 70dB SPL range with 10 dB SPL steps, and 2 – 4 s ISI. Sound intensity and ISI were presented pseudo-randomly. Behavioral recording sessions lasted 1 hr. A water pump (AL-1000, World Precision Instruments) and sound presentation were controlled by a custom-written LabVIEW program (National Instruments). Licking sensor (PMU24, Panasonic) information was relayed through NI-DAQ (NI PCI-6221, National Instruments).

### Histological analysis

Mice were injected intraperitoneally with a mixture of pentobarbital and lidocaine and perfused transcardially with PBS followed by 4% paraformaldehyde (PFA) in 0.1M PBS. Brain tissue was removed and stored in the same fixative overnight at 4°C, then transferred into 30% sucrose solution for at least 2 days. The tissue was cut into 80 µm coronal sections using a microtome (SM2010R, Leica), stained with DAPI, and observed under an epifluorescent upright microscope (Eclipse E600, Nikon). In adult mouse brain slices, post electrophysiological recordings, DiI/CM-DiI signals were also observed to evaluate the probe track.

### Data analysis

#### Histological evaluation of superficial cortical layers

Width of cortical layers in P2 and P21 mice was measured using Fiji ImageJ on DAPI images taken with 4× magnification. Portion of superficial layers in the cortex was calculated using the formula: *W_L2-4_*/*W_L2-6_*, where *W_L2-4_* and *W_L2-6_* were the width of L2-4 and L2-6, respectively. Measurements of 3 adjacent areas were taken on each brain slice, one brain slice per mouse, and the mean of their ratios was compared to other brain slices. All measurements were done blindly to the treatment and grouped afterwards. In P2 mice, the measurements were taken in the frontal, somatosensory area, while at P21 the measurements were taken in the auditory cortical area.

#### Behavioral data

In each session, mean reaction time and detectability index (*d’*) were calculated for all sound intensities. *d’* was calculated using the formula: *d’* = *Z*(hit rate) - *Z*(false alarm rate), where Z is the inverse of the standard normal cumulative distribution function. The detection threshold of each training session was estimated based on cross-validated logistic regression analysis. A logistic binomial model of the probability of hit as a function of sound intensity was created using MATLAB *fitglm* function. The detection threshold was defined as the sound intensity which reaches 50% hit. This process was repeated with a randomly chosen half of trials and the remaining half, and the threshold was calculated as the mean of them.

#### Spike sorting

All electrophysiological data analysis was performed offline. In silicon probe recordings, spike sorting was performed with Kilosort or Kilosort2 (https://github.com/MouseLand/Kilosort), followed by manual curation with phy2 (https://github.com/cortex-lab/phy). Quality of each cluster was assessed by their Mahalanobis (isolation) distance (Schmitzer-Torbert et al., 2005). In all further assessments, we included only the single units with an isolation distance ≥ 20 and an overall spike rate >0.1 Hz.

In Neuropixels recordings, spike sorting was performed with Pykilosort (https://github.com/MouseLand/pykilosort). Quality of the clusters recognized as “good” by Pykilosort was assessed through additional metrics: sliding refractory period violation and a noise cutoff estimate (Banga et al., 2022). Briefly, the first metric estimated whether each cluster is contaminated by refractory period violations without assuming the refractory period duration. The criterion was set as clusters with <10% contamination. The second metric estimated to what extent spikes were detected without cutting off by the detection threshold. Computing the amplitude distribution of detected spikes, this metric computed how many standard deviations (SDs) of the low bin falls outside the mean number of spikes in the high quantile. The criterion was set as follows: the value was <5 SDs and the value of the lowest bin was <10% of the value of the highest bin. Only the “good” clusters passing all thresholds were considered as single units. Out of these units, those with an overall spike rate >0.1 Hz were included in further evaluations.

#### Cortical cell classification

Cortical cells were further classified based on their spike waveforms (Yague et al., 2017). We set the threshold of the trough-to-peak duration to 0.5 ms to distinguish between broad-spiking (BS) and narrow-spiking (NS) cells. In the analysis of electrophysiological data from task-performing mice, we have only included the recordings of sessions where the mice passed the behavioral performance threshold (difference between the false alarm rate and 70 dB hit rate higher than 40%).

#### Probe track estimation

To estimate which channels belong to which brain region in Neuropixels recordings, we assessed the overall multi-unit activity on each probe channel and cross-compared it with a 3D histological evaluation of the probe’s trajectory made by SHARP-Track (https://github.com/cortex-lab/allenCCF/tree/master/SHARP-Track (Shamash et al., 2018)). Combining SHARP-Track information with known brain-area specific activity allowed us to determine which recording channels were located in the auditory cortex and which were in the thalamus. Because SHARP-Track borders are not exact, as they are affected by tissue distortion and tearing during slice fixation, we have calculated the median absolute deviation between our electrophysiological and histological borders. On average, this deviation was 69 μm, which is comparable with a previous report (Steinmetz et al., 2019).

To define the MGB borders, we took the following two-step estimation approach: (1) the estimation of the number of channels within MGB based on histological analysis, (2) the refinement of the channel positions based on auditory evoked responses. More specifically, by taking the output of the SHARP-Track analysis described above, we estimated the length of MGB passed by the Neuropixels probe. This length was converted to a corresponding number of electrode channels based on the spacing between electrodes. Secondly, we determined the latency of auditory evoked responses across all thalamic channels. For each channel, cell activity was split into 1 ms bins, filtered using 5 ms Gaussian kernel and normalized by evoked response threshold (2.8 SD from baseline mean, taken from 100 ms before the sound onset across trials). Therefore, all values above 1 were considered a significantly evoked response and latency was determined as the first occurrence of such response. Since most MGB neurons have a short latency of auditory stimuli (Anderson and Linden, 2011), we identified a cluster of channels exhibiting short response latency (<20 ms), and refined the MGB channel positions while maintaining the number of channels estimated by the first step.

#### Cortical laminar estimation based on current source density (CSD) analysis

In silicon probe recordings, broadband signals from each channel were lowpass filtered (800 Hz), downsampled (1 kHz) and spatially smoothed across one top and one bottom neighboring channel. In Neuropixels probe recordings, local filed potential (LFP) signals were lowpass filtered (100 Hz), downsampled (1 kHz) and spatially smoothed across ±5 channels. Event-related potentials (ERP) were then calculated and filtered using lowpass and highpass Butterworth filter, and CSD was calculated using the formula: CSD = ((*Va*+*Vb*)-2*Vo*)/*d*^2^, where *Vo* is the observed channel, *Va* and *Vb* are its neighboring channels and *d* is the distance between channels (mm). CSD sink channel was set as the top border of input layer (L_in_), which was then counted as depth 0, while the borders of deep layer 1 (L_d1_; putative L5) and deep layer 2 (L_d2_; putative L6) were determined based on layer thickness from previous publications (Cooke et al., 2018, Ji et al., 2016). Thus, L_d1_ top border was set at –150 μm and L_d2_ top border at –400 μm. Everything above 0 μm was considered as superficial layers (L_sup_; putative L2/3). Out of 22 passive listening Neuropixels recordings, 7 recordings (1 from control and 6 from treated mice) were excluded from further laminar and LFP analysis due to excessive noise levels in LFP channels.

#### Spontaneous activity analysis

In passive listening mice, the spontaneous firing rate of each cell was calculated during a 5 min silent period recorded immediately after the noise presentation period. In the same silent period, Welch’s power spectral density was calculated for five frequency bands (delta [1-4 Hz], theta [4-8 Hz], alpha [8-12 Hz], beta [15-30 Hz] and low gamma [30-48 Hz]) for each channel. Signals from the silicon probes were first extracted with a lowpass filter (800 Hz) and then downsampled to 1 kHz. Only the channels from one (middle) shank were taken into further analysis. For Neuropixels recordings, only one row of channels was used in LFP analysis. LFP signals were lowpass filtered (50 Hz) and spatially smoothed across ±2 channels before power spectral density calculation. For estimation of mean power per cortical layer, CSD results were used to split the channels into layers, and the mean relative power per layer was calculated for each recording. For cortical and hippocampal mean power estimation, the power across all channels in the auditory cortex or hippocampus was calculated, and their mean per recording was used in group comparisons. In task-performing mice, spontaneous activity was estimated from 0.5 s periods before the sound onset.

#### Auditory evoked neuronal activity

For peri-stimulus time histogram (PSTH), average activity during noise presentation of different intensities was calculated in every 1 ms bin across stimulus conditions. For statistical analysis of evoked neuronal activity, spike counts during the 50 ms time window from the sound onset were used.

#### Cross-correlation analysis

Monosynaptic connections assessment was done as described elsewhere (Fujisawa et al., 2008). After computing the original cross-correlograms (CCGs), 1000 surrogate CCGs were created to assess statistical significance. To create each surrogate CCG, spikes were randomly jittered ±3 ms and a CCG was calculated as a surrogate. Based on these surrogate CCGs, upper and lower bounds of confidence limit (1%; *p* = 0.01) were determined. When the original CCG values passed the confidence limit within the [-3:0 ms, 0:3 ms] time window, the interaction between cells was considered significant (*p* < 0.01). If the values passed the upper confidence limit, the interaction between neurons was considered as excitatory. Otherwise, if the values passed the lower confidence limit, the interaction was labelled as inhibitory.

### Statistical analysis

All statistical analysis was done using MATLAB. The effect was considered significant at *p* < 0.05. In Figures 1C, 1D, 1E, 1F, 4A and 4C two sample *t*-test was performed. In Figures 2D and 2E a two-way ANOVA was performed. Rank sum test was done in Figures 2D (inset), 3E, 3F, 6B and 7C. Rank sum test with Bonferroni correction was performed in Figures 5C, 5D, 5E, 5F, 6C, 6D, 6E, 6F, 7D, 8C and 8D. To estimate effect size, Hedge’s *g* was calculated in Figure 4B, using a MATLAB toolbox (Hentschke and Stuttgen, 2011).

